# The temporal evolution of cancer hallmarks

**DOI:** 10.1101/2024.01.21.576566

**Authors:** Lucie Gourmet, Daniele Ramazzoti, Parag Mallick, Simon Walker-Samuel, Luis Zapata

**Affiliations:** Centre for Computational Medicine, University College London, London, United Kingdom; Department of Medicine and Surgery, University of Milano-Bicocca, Monza, Italy; Canary Center for Cancer Early Detection, Stanford University, Palo Alto, United States; Centre for Evolution and Cancer, Institute of Cancer Research, London, UK

**Author notes:** **Correspondence:** Dr. Luis Zapata.

**Keywords:** cancer, hallmark, evolution, mutations, genome instability, immune evasion

## Abstract

Cancer hallmarks describe key physiological characteristics that distinguish cancers from normal tissues. The temporal order in which these hallmarks appear during cancer pathogenesis is of interest from both evolutionary and clinical perspectives but has not been investigated before. Here, we order hallmarks based on the allele frequency and selective advantage of mutations in cancer hallmark genes across >10k untreated primary tumors and >8K healthy tissues. Using this novel approach, we identified a common evolutionary trajectory for 27 of 32 cancer types with genomic instability as the first and immune evasion as the last hallmark. We demonstrated widespread positive selection in cancer and strong negative selection in normal tissues for all hallmarks. Notable exceptions to the hallmark ordering in tumours were melanomas (uveal and skin) suggesting that strong environmental factors could disrupt common evolutionary paths. Clustering of hallmark trajectories across patients revealed 2 clusters defined by early or late genomic instability, with differential prognosis. Our study is the first to identify the temporal order of cancer hallmarks during tumorigenesis and demonstrate a prognostic value that could be exploited for early detection and risk stratification across multiple cancer types.

## Main text

Cancer is an evolutionary process described as the sequential acquisition of genomic alterations affecting cellular hallmarks under positive selection (1). Hanahan and Weinberg first identified six cancer hallmarks (2), later extended to ten (3,4): self-sufficiency in growth signals, insensitivity to growth-inhibitory signals, evasion of programmed cell death, limitless replicative potential, sustained angiogenesis, metastasis, genome instability, tumour-promoting inflammation, avoiding immune destruction, and deregulating cellular energetics. These hallmarks are established functions of tumour cells identified over several decades exploiting a wide array of scientific and technological approaches. Advanced imaging modalities, including magnetic resonance imaging (MRI), computed tomography (CT), ultrasonography, and positron emission tomography (PET) have revealed morphological changes of cancer cells associated to hallmarks (5). Intravital microscopy has shown the interactions between tumor vasculature, immune interactions, and cellular proliferation (6). CRISPR-based methodologies have uncovered cancer hallmarks by monitoring genome instability (7) and epigenetic editing (8). Genomic screening has enabled the identification of genes implicated in metastasis and metabolic regulation (9,10) and genomic analysis of large cancer datasets has linked specific genes to distinct cancer hallmarks (11,12).

Modern sequencing technologies allow the estimation of cell proportions carrying a given somatic mutation (i.e., using variant allele frequency or VAF). In parallel, computational methods and the vast number of genomic databases enable the annotation of the functional impact of these mutations. Recently, VAF and the ratio of non-synonymous to synonymous mutations (dN/dS) have been used to explore various facets of cancer evolution. These include the identification of neutral (13), positive (14,15) and negative selection during tumorigenesis (16), the impact of immune selection against neoantigens (17,18), and the observation of positive selection in healthy tissues (19–21). These metrics have further been used to trace the phylogenetic relationship and intratumor heterogeneity among various cancers (22,23) but only few have focused on understanding the emergence of tumorigenic properties in somatic tissues (24). Also, cancer hallmarks have been studied as ecologically driven (25) or as spatially distributed phenotypes (26). However, the temporal order in which they appear in the pathogenesis of individual tumours is unknown.

Here we combined two evolutionary metrics, variant allele frequency (VAF) and the ratio of non-synonymous to synonymous mutations (dN/dS), to establish the temporal order and the selective advantage of each cancer hallmark across multiple tumor types and in healthy tissues.

## Results

### A common temporal hallmark ordering across multiple tumor types

To determine the temporal order of cancer hallmarks among 32 tumor types (Supplementary Table 1), we used the variant allele frequency of nonsynonymous somatic point mutations (VAF, Fig. 1a) in genes associated with each hallmark as a surrogate of time. Among the gene lists for in hallmarks (Fig 1b, Supplementary Table 2), angiogenesis harboured the largest proportion of shared genes with any other (100%, 478/478), followed by growth (93%, 487/522) and metastasis (92%, 982/1072). The hallmarks with the fewest shared genes were metabolism (59%, 251/429) and genome instability (41% 87/211). Intriguingly, less than half of the known cancer driver genes were associated with any specific hallmark (42%, 152/365) and our previous list of escape genes (18) shared almost 70%. Among the genes most represented in the ten hallmarks we found AKT, Ras, PIK3 and MAPK gene families (9 out of 10 hallmarks), and also other well characterised genes such as BRAF (7/10) and TP53 (8/10).

**Figure 1.**
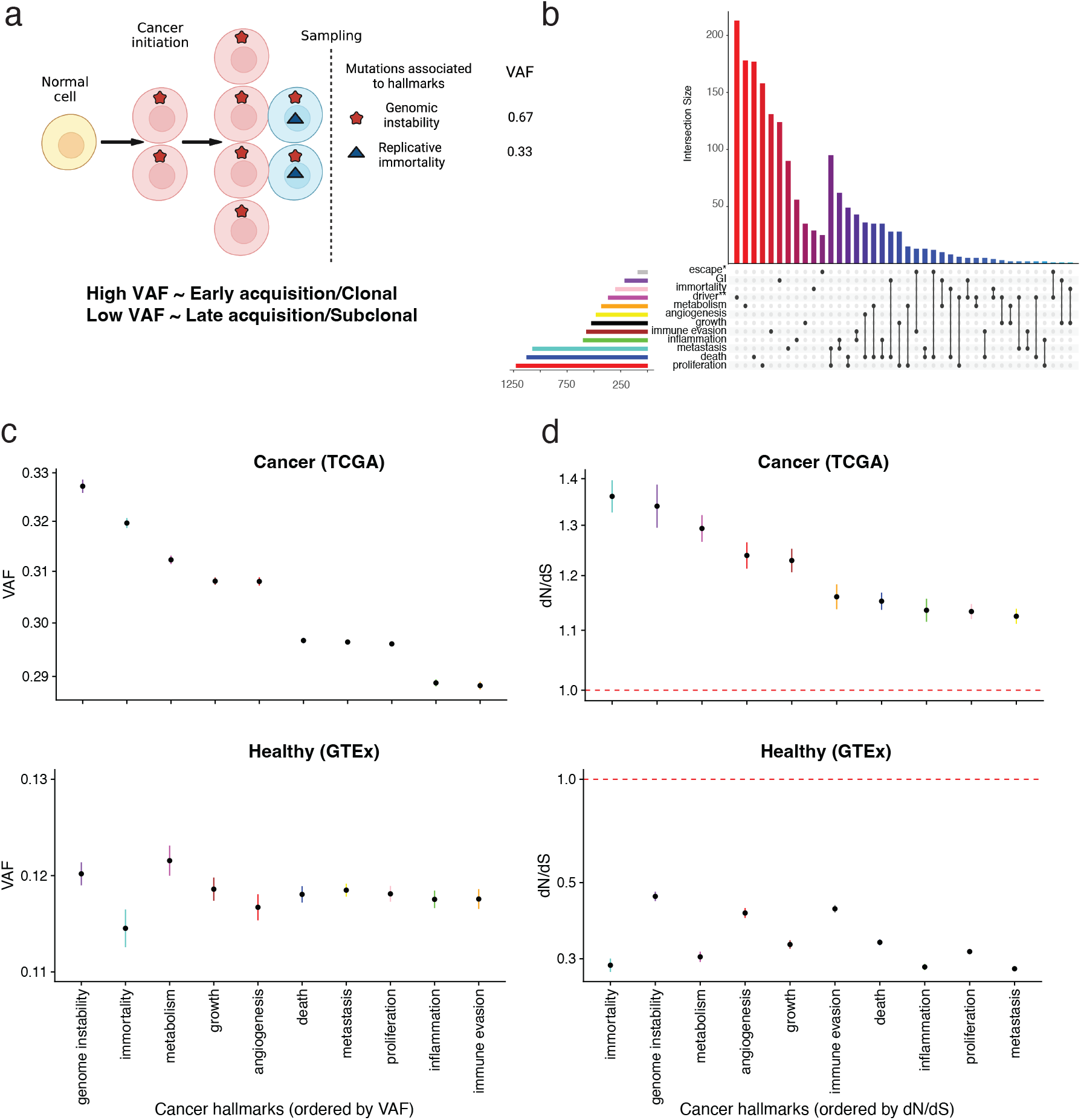
Temporal Ordering of Cancer Hallmarks Using VAF and dN/dS in Cancer and Normal Tissues. (a) The mean VAF of somatic mutations in various cancer hallmark-associated genes is estimated as a proxy for timing; hallmarks selected early will exhibit higher allele frequencies. (b) Overlap among genes associated with cancer hallmarks taken from Zhang et al 2020. Two other list of genes were used as controls, escape genes (*) from Zapata et al 2023, and driver genes (**) from Martincorena et al 2017. (c) Mean Variant Allele Frequency (VAF) for each cancer hallmark, derived from mutations in all primary tumors in the TCGA database and normal tissue from GTEx data. A higher VAF suggests an earlier appearance of the hallmark, thereby allowing inference of their relative ordering. Data points represent the mean VAF, and error bars indicate the standard error. (d) Mean dN/dS value associated with each cancer hallmark, derived from mutations in all primary tumors in the TCGA database and normal tissue from GTEX data. All cancers exhibited dN/dS > 1, indicative of positive selection for these hallmarks. In contrast, normal tissues showed dN/dS < 1, signifying negative selection in healthy tissues.

We calculated the mean and standard error of VAF for all cancers together (pan-cancer) and for all non-cancerous samples. As synonymous events do not confer a selective advantage to the cells, we also controlled for random drift by repeating this analysis using only synonymous mutations (Supplementary Fig. 1). In the pan-cancer analysis (Fig. 1c), we found that genomic instability (VAF = 0.3272 ± 0.001) and replicative immortality (VAF = 0.3196 ± 0.001) occurred first in cancer development, suggesting that these altered functions are required at an early stage to promote the accumulation of subsequent mutations and to outcompete normal cells by avoiding telomere attrition, an effect known as the “Hayflick limit” (27). The following hallmarks were metabolism (VAF = 0.3123 ± 0.0008), evading growth suppressors (VAF = 0.3081 ± 0.0007), angiogenesis (VAF = 0.3080 ± 0.0008), resisting cell death (VAF = 0.2967 ± 0.0005), metastasis (VAF = 0.2964 ± 0.0004), and proliferative signalling (VAF = 0.2961 ± 0.0004). Cancer cells deregulate their metabolism possibly due to increasing pressure on strategies to metabolize glucose under hypoxic conditions, also known as the Warburg effect (28). As the tumor grows, angiogenesis will support the rapid growth of tumor cells and alleviate hypoxia (29), and then resistance to cell death (30) will avoid intrinsic (from aberrant or dysregulated pathways) and extrinsic death signals (from the tumor microenvironment). Sustaining proliferative signals is acquired in the middle of tumorigenesis, possibly thanks to the intrinsic cell cycle checkpoint mechanisms developed to avoid self-destruction. The last two hallmarks, inflammation (VAF = 0.2888 ± 0.0007) and immune evasion (VAF = 0.2883 ± 0.0007), are involved in the adaptation of tumor clones to the immune microenvironment and are often co-occurring, despite having less than 10% of overlapping genes (Fig. 1b). In contrast, healthy tissues display no clear order with overlapping VAF estimates for 6 out of 10 hallmarks, indicating that the previous order was cancer-specific (Fig. 1c). We must emphasize that this understanding of cancer pathogenesis was reconstructed from VAFs of point mutations without including gene expression or copy number alterations in the genes involved in each hallmark. However, when we include only diploid tumors in our analysis the results are similar (Supplementary Fig. 2). In controls using only neutral genes, we showed that VAF is significantly lower compared to hallmark genes (Supplementary Fig. 3).

Next, to determine which hallmarks held the most significant fitness advantage, we computed the ratio of nonsynonymous to synonymous mutations (dN/dS). In cancer, all hallmarks exhibited a dN/dS greater than 1, consistent with positive selection (Fig. 1d). This finding contrasted starkly with normal tissues, where all hallmarks demonstrated a dN/dS less than 1, indicative of pervasive negative selection against tumorigenic properties in healthy tissues. Genomic instability and immortality hallmarks were subject to the strongest positive selection in cancer (genomic instability: 1.340, 95% confidence interval [CI] = [1.295, 1.387]; immortality: 1.361, CI = [1.327, 1.396]), while in non-malignant tissues displayed dN/dS less than one (immortality: 0.287, CI = [0.275, 0.300]; genomic instability: 0.456, CI = [0.442, 0.471]). After these two hallmarks, we identified metabolism, angiogenesis, growth evasion, death, inflammation, proliferation, and finally metastasis.

We further investigated the sequence of hallmarks by integrating both VAF and dN/dS into a single metric (Methods). However, the results were inconsistent with our previous order, with metastasis as the first hallmark, suggesting this strategy was not appropriate for determining the temporal order of hallmarks (Supplementary Fig. 4). To avoid including genes without a clear impact on tumorigenesis, we focused only on genes under significant positive selection for each hallmark. We confirmed that ‘genomic instability’ was first, while ‘inflammation’ and ‘immune evasion’ were the last two (Supplementary Fig. 5). Interestingly, most of genes under significant positive selection are known drivers of tumorigenesis, including ATM, CASP8, TP53 and PTEN in genome instability, and PIK3R1 and HLA genes for inflammation and evasion (Supplementary Fig. 6). We also repeated the analysis using VAF corrected by ploidy (Supplementary Fig. 7) with no observed differences, but when using cancer cell fraction (CCF), which includes information from copy number and ploidy status, we found that genomic instability fell to a middle position, while immune evasion remained the last (Supplementary Fig. 8). Finally, we also explored the impact of removing TP53, the most commonly gene mutated in cancer, from all hallmarks and found that genome instability became last, while most other hallmarks were not affected (Supplementary Fig. 9), suggesting that early acquisition of genome instability is predominantly driven by TP53.

### Environmental pressures influence inter-tumor hallmark heterogeneity

To determine inter-tumor heterogeneity on the temporal order of cancer hallmarks, we computed the hallmark mean VAF ranked for each tumor (from 1 to 10, Supplementary table 3 and 4), and computed their Pearson correlation for all versus all (Fig. 2a). Strikingly, the ordering of hallmark appearance was consistent across tumor types, with most tumor types following a common trajectory, whereas only five showed deviations (e.g., uveal melanoma, thymoma, thyroid carcinoma, skin cutaneous melanoma, pheochromocytoma and paraganglioma). When looking at only the significant correlations corrected by multiple testing, nine tumor types did not share a common trajectory with the rest (Supplementary Fig. 10). In addition, we observed that evolutionary trajectories were less consistent between cancer types when using CCF (Supplementary Fig. 11), dN/dS-based rank (Supplementary Fig. 12), or when removing TP53 from all analysis (Supplementary Fig. 13). However, when adapting a recently developed algorithm, ASCETIC, to infer evolutionary trajectories based on genes to gene groups (24), we found that most tumor types followed a similar order of cancer hallmark acquisition (Fig 2b).

**Figure 2.**
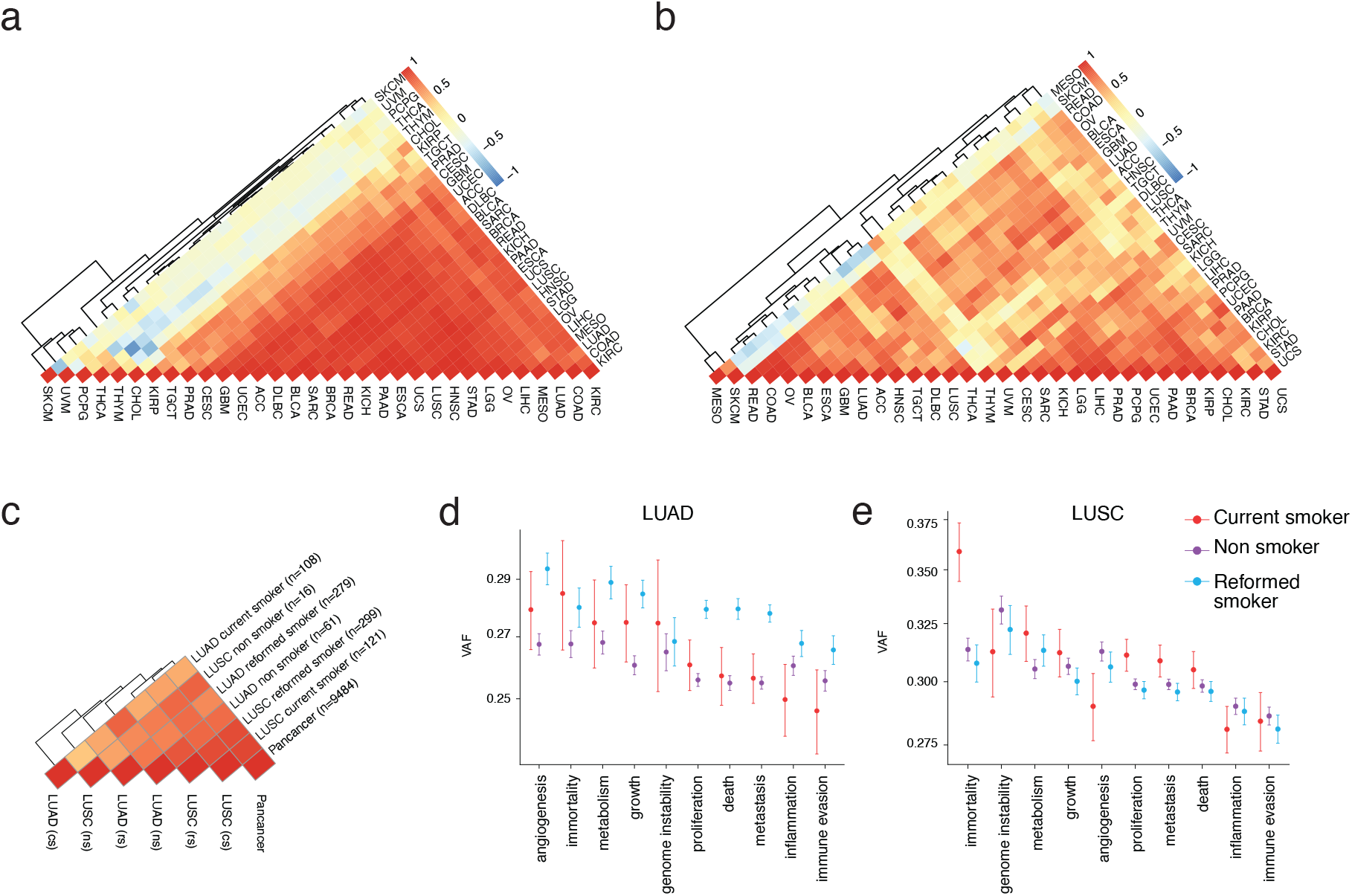
Comparison of Cancer Hallmarks order between Subtypes of Cancer. a) Hierarchical clustering of spearman correlation values of the rank order according to VAF for 32 cancer subtypes. b) Hierarchical clustering of spearman correlation values of the rank order obtained using ASCETIC algorithm. c) Hierarchical clustering of spearman correlation values of the rank order according to VAF for Lung tumor separated by smoking status. VAF values associated with each cancer hallmark for smokers, former smokers, and non-smokers in (d) Lung Adenocarcinoma (LUAD) and (e) Lung Squamous Cell Carcinoma (LUSC).

As skin and uveal melanomas consistently exhibited differences from other tumor types and are well-documented to be influenced by UV light exposure, we hypothesized that specific environmental factors could affect the evolutionary trajectories of cancers. To investigate this further, we focused on two types of tumors with available data on smoking status: lung squamous cell carcinoma (LUSC) and lung adenocarcinoma (LUAD). The pair with the most divergent trajectories were LUAD current smokers and LUSC non-smokers suggesting that cell of origin and environmental factors influence the temporal order of cancer hallmarks. We found that LUSC patients overall had an evolutionary trajectory similar to pan-cancer (R_s_ = 0.94) (Fig 2c). Lung adenocarcinoma (LUAD) patients with a smoking history had a significantly higher overall VAF compared to non-smokers and current smokers across all hallmarks except for genome instability (Fig. 2d), suggesting a clonal sweep after smoke exposure driven by modification of metabolism, growth capacity and angiogenesis (the top 3 ranked hallmark in this group). Smokers, non-smokers, and reformed smokers with LUSC have similar VAF values for seven out 10 hallmarks. LUSC current smokers had higher VAF in immortality, proliferation, and metastasis (Fig. 2e). Interestingly, inflammation and immune evasion displayed low ranks (last) compared to the other hallmarks in all LUSC groups, and in all but non-smokers of LUAD where they were no longer the last, suggesting that smoking might be less associated to immune evasion than previously reported.

To compare if the temporal order inferred from VAF and dN/dS was consistent, we determined their rank-based correlation. While VAF and dN/dS-based ordering were significantly correlated in pancancer analyses (Spearman’ s rank correlation = 0.806, p = 0.008) there was no correlation observed in normal tissues (Spearman’ s rank correlation = 0.103, p = 0.785), and the correlation was less clear when comparing tumor types (Supplementary Fig. 14). Moreover, when comparing the temporal order between cancer and normal tissue we did not find a significant correlation when using dN/dS (Spearman’ s rank correlation = 0.382, p = 0.279) or VAF (Spearman’ s rank correlation = 0.309, p = 0.387).

### Common routes of cancer hallmarks determine cancer prognosis

To elucidate the temporal order of cancer hallmarks in individual patients, we ranked each hallmark from one to ten, based on the mean variant allele frequency (VAF) of mutations in the genes associated with each hallmark (Supplementary table 5 and 6). Hallmark frequencies for each individual yielded results consistent with our pan-cancer and inter-tumor analyses (Fig. 3a). The hallmark of genomic instability showed a dichotomous pattern predominantly being either the first or last event (ranked 1 or 10), which depended mostly on the presence of mutations in TP53 (Supplementary Fig. 15). Half of individuals with GI hallmark mutated were due to a TP53 mutation (Supplementary Fig. 16). Early acquisition trends were noted for hallmarks such as angiogenesis, cellular growth, immortality, and metabolic processes, with their prevalence diminishing in later stages. In contrast, hallmarks related to cell death, proliferation, and metastasis showed a pattern of later acquisition, though not necessarily as the latest events in tumor evolution. Inflammation and immune evasion displayed similar late trajectories, suggesting that both hallmarks generally manifest in the final stages of the tumour’s evolutionary course, even after the metastatic abilities have been acquired.

To identify differences between individual patients, we clustered all samples based on the ranks (Fig. 3b). Principal component analysis revealed that genomic instability (GI), immune evasion, and inflammation (Fig. 3c) were the primary features distinguishing the patient cohort into two distinct classes (Fig. 3d). These classes were prognostic of overall survival, progression-free survival, and disease-free survival rates (Fig. 3e). Patients with an early acquisition of the genomic instability hallmark typically had poorer outcomes (Cluster 2), while a later acquisition was associated with better prognoses. A cross-tumor type analysis showed an even distribution of patients across the two clusters (Fig. 3f), with some exceptions in specific tumor types such as breast (BRCA), head and neck squamous cell carcinoma (HNSC), lung squamous cell carcinoma (LUSC), and ovarian (OV) cancer. In addition, we used the ASCETIC algorithm to clusters patients and found a prognostic difference when 6 clusters were defined (Supplementary Fig. 17). Next, we compared the proportion of individuals with each hallmark at that rank (Supplementary Fig. 18). The top three hallmarks at rank 1 were genome instability, metabolism, and immortality, while at rank 10, they were genome instability, inflammation, and immune escape, corroborating our previous results. To control for mutation biases in hallmark ordering or selected gene lists, we randomized the labels of all genes associated with each hallmark, recomputed the VAF-based ranks, and found that hallmarks were evenly distributed across all ranks (Supplementary Fig. 19).

**Figure 3.**
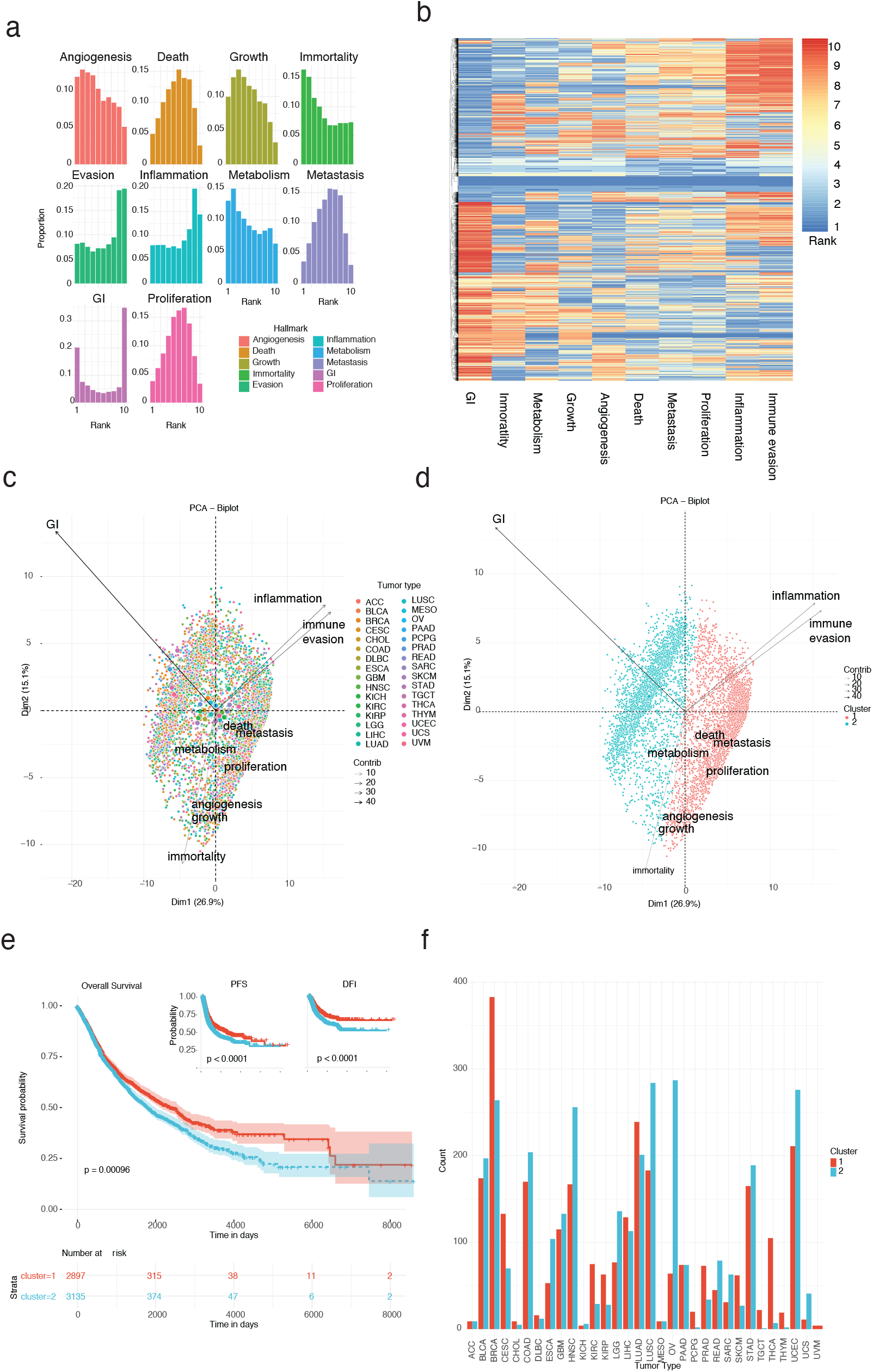
Patient trajectories, tumor type heterogeneity stability on patient trajectories. a) Proportion of individuals with a certain rank for each hallmark. B) Hierarchical clustering of all patients based on the hallmark ranks. C) PCA plot of all patients based on their hallmark rank. D) Same as C but labelling patients based on the cluster assigned. E) Overall survival of patients based on two clusters. F) Proportion of individuals per tumor type in each cluster.

## Discussion

Cancer hallmarks are recognized as defining attributes of tumors, identified through extensive research over decades. The order in which they are acquired during tumor pathogenesis has remained unclear. Leveraging a unified genetic framework, our study reveals the temporal order of these hallmarks across various tumor types for the first time. By associating hallmark characteristics with specific genetic mutations and analysing variant allele frequency (VAF) along with the nonsynonymous-to-synonymous mutation ratio (dN/dS), we discern the relative timing and selection pressures for each hallmark. This analysis spans over 10,000 samples from 32 cancer types and contrasts with 8,000 healthy tissue samples.

Synthesizing data across cancers, we observe a general order of cancer hallmarks trough cancer evolution with early genomic instability and replicative immortality in primary tumors, and immune evasion, metastasis, and inflammation hallmarks typically arising later. The occurrence of each hallmark is directly correlated with the degree of positive selection it experiences in cancer, and contrast with negative selection in healthy tissues. Notably, excluding TP53 mutations from our analysis shifts genomic instability to the end of the hallmark sequence, suggesting its pivotal role on enabling cancer via increased disruption of mutation accumulation.

Modelling cancer hallmark evolution has seen various approaches, from the agent based CancerSim (31) modelling six hallmarks to the more recent tugHall, which employs an approximate Bayesian computation against TCGA colorectal cancer data (32). These models, however, often adopt a reductionist gene-to-hallmark mapping, not fully representing the complexity of cancer genetics where genes like TP53 influence multiple hallmarks.

By understanding the order of cancer hallmarks, we can target specific hallmarks at specific stages. Knowing cancer hallmarks evolutionary order enables early cancer intervention and improves personalised medicine. Moreover, hallmark ordering provides information for prognosis: different evolutionary trajectories are likely associated with different survival outcomes. Understanding which hallmarks occur first/last will change our approach to cancer experiments. If genome instability is required early for carcinogenesis, cancer models will need to account for that (for example by knocking out TP53 to create precursor lesions instead of focussing on other hallmark related genes). It is experimentally challenging to investigate the hallmark ordering, but our bioinformatics study can shed light on this challenge.

The established temporal sequence of cancer hallmarks enriches our understanding of tumor biology, enabling a novel patient stratification based on genetic evolution. This stratification could transform prognosis and treatment, particularly by anticipating and countering immune evasion. Our work lays the groundwork for this transformative approach in oncology, offering a genetic-based framework to navigate the complex landscape of cancer evolution.

## Methods

### Data Acquisition and Pre-processing

Cancer hallmark genes were obtained from the study by Zhang et al., with each hallmark being associated with a specific gene set (21). These gene sets were filtered to match the current HUGO Gene Nomenclature Committee (HGNC) database. Two additional gene categories, escape and driver, were compiled to provide a comparative reference for the immune evasion hallmark (Zapata et al., 2023) and a collection of cancer driver genes (Martincorena et al., 2017), respectively. We created an upset plot to illustrate the overlap between the hallmark categories and the size of the gene lists using the r package UpSetR v1.4.0.

Each hallmark was given a shorthand name:

- Sustained proliferative signalling: proliferation
- Evading growth suppressors: growth
- Resisting cell death: death
- Inducing angiogenesis: angiogenesis
- Activation of invasion and metastasis: metastasis
- Enabling replicative immortality: immortality
- Avoiding immune destruction: immune evasion
- De-regulation of cellular energetics: metabolism
- Genome instability and mutation: genome instability
- Tumour promoting inflammation: inflammation

We used data from TCGA (https://portal.gdc.cancer.gov/) and GTEX (normal tissue) to get the variant allele frequency (VAF) value for every mutation (33). TCGA data were annotated using the annotation step from dNdScv, excluding metastatic samples and duplicated data, retaining only samples from primary tumors. The TCGA pan-cancer dataset comprises 2,626,225 mutations across 9,484 patients from 32 cancer types (https://gdc.cancer.gov/resources-tcga-users/tcga-code-tables/tcga-study-abbreviations). The GTEX pantissue dataset includes 999,203 mutations across 7,584 patients from 16 tissue types: Adipose Subcutaneous, Adipose Visceral Omentum, adrenal gland, aorta, coronary artery, Artery Tibial, Brain Caudate basal ganglia, Brain Cortex, Brain Frontal Cortex BA9, Brain Hippocampus, Brain Hypothalamus, Brain Nucleus accumbens basal ganglia, Brain Putamen basal ganglia, Breast Mammary Tissue, Colon Sigmoid, Colon Transverse.

### VAF and dN/dS calculation

VAF is a measure of the number of copies of a specific mutated allele, relative to the total number of alleles (4), which is widely used to measure the timing of acquisition of the mutated allele during tumor pathogenesis(34,35). To determine the order of hallmarks using VAF, we filtered the mutations by impact (Missense, Nonsense, Essential Splice, and Stop loss) to retain only nonsynonymous mutations. We then generated a data frame for each hallmark containing patient ID, gene name, and VAF. For each hallmark we calculated the mean VAF and standard error. These were plotted and ordered from high VAF (early) to low VAF (late) with standard errors represented as error bars, using the ggplot2 package in R.

We run dNdScv R package (version 0.0.1.0) to examine the selection pressures on each cancer hallmark. We used the genelist option providing the list of genes associated with each hallmark (15). We obtain the global dN/dS output (dN/dS calculated using all mutations from the gene list) from dNdScv and plotted such value along with the upper (dndshigh) and lower (dndslow) 95% confidence intervals. The hallmarks were ordered based on higher dN/dS (strong positive selection if > 1) to lower dN/dS.

To create correlation heatmaps between cancer types, we calculated the mean VAF of each of the 10 hallmarks for every cancer type separately. We then determined the Spearman correlation between tumor types and used the pheatmap R package for visualization. The same steps were followed to produce a heatmap for dN/dS and to assess the relationship between dN/dS and VAF.

To normalize for biases on tumor purity between tumor types, we determined the rank for each hallmark in each patient based on the mean VAF. We then calculated the proportion that each hallmark was first, second until the last 10, filtering for patients that did not have information in at least 8 hallmarks. We plotted the distribution at each rank with and without TP53. We plotted the oncoprint for all patients for genome instability using the library MAFtools.

### Alternative metrics for timing VAF and estimating dN/dS

To determine VAF distributions in other known gene sets, we estimated mean VAF from: 1) “driver,” comprising 365 cancer driver genes known to be under positive selection, 2) “escape,” incorporating a published list of genes associated with immune evasion, and 3) “neutral”, where we sampled 200 random genes not included in any hallmark to create neutral categories. In addition, we calculated a corrected version of VAF, called Cancer Cell Fraction (CCF), to account for tumor ploidy and purity using the formula CCF = (VAF/m*p) * (p*N + 2*(1-p)) where VAF is the allele frequency of the alternative allele, m is the inferred multiplicity of the mutations, p is the sample purity, and N is the local copy number, as described in a previous study (36). Some CCF values exceeded 1, so they were excluded from our dataset to ensure the validity of our results. We also performed the VAF analysis ordering the hallmarks only using tumors where ploidy was 2 (diploid).

### Clustering analysis

Initially, the raw dataset, which included median VAF for 10 hallmarks from all TCGA samples, underwent pre-processing to ensure suitability for analysis. This involved checking for missing values and outliers, with the distribution of values assessed visually through boxplots and histograms using ggplot2 in R. Feature scaling was then applied to normalize the data, as clustering algorithms like k-means are sensitive to variable scales. This normalization was achieved using the scale function in R, standardizing each variable to have a mean of 0 and a standard deviation of 1.

Principal Component Analysis (PCA) was applied for dimensionality reduction using the prcomp function in base R. This step summarized the key features of our data in fewer dimensions, facilitating more efficient clustering. To identify the optimal number of clusters for k-means, the Elbow method was used. We plotted the total within-cluster sum of squares (WSS) against a range of potential cluster numbers using the fviz_nbclust function from the factorextra package in R. The ‘elbow point,’ where the decrease in WSS diminished noticeably, indicated the optimal number of clusters.

K-means clustering was then executed with the kmeans function in R, after determining the optimal number of clusters. The cluster assignments for each sample were recorded. The quality of our clustering was assessed using the silhouette method, which evaluates how similar a sample is to its own cluster relative to other clusters. We computed the silhouette scores for each sample using the silhouette function from the cluster package in R. The average silhouette width was calculated to assess the overall quality of the clustering, with higher average silhouette widths indicating better-defined clustering. Finally, the clusters were visualized in a two-dimensional PCA space using a scatterplot created with ggplot2, with different colours representing different clusters. This allowed us to visually inspect the spatial segregation of the clusters and gain an intuitive understanding of the sample groupings.

### Randomisation analysis

To ensure the validity of our results, we investigated the distribution of the mean VAF of random genes (called neutral if not associated to any hallmark) across nonsynonymous mutations. We created 10 categories each composed of 473 genes (473 being the mean number of genes associated to hallmarks) to calculate the mean VAF of these categories. This step was performed 100 times on hallmarks genes only and on neutral genes only to compare the two distributions in a histogram. Besides, to validate the ordering of the hallmarks we created pseudo hallmarks (categories of the same size as the original hallmarks but sampling from neutral genes). We then plotted the frequency of ranks for each hallmark for these pseudo hallmarks and compared it to the frequency of our actual results.

### ASCETIC analysis

Hallmarks data used as input for ASCETIC were derived from genes associated with Hallmarks, as described previously. Specifically, for each Hallmark and sample, a binary matrix was generated, indicating 1 if the Hallmark was observed in the sample and 0 otherwise. Additionally, mean CCF and VAF data were employed to estimate the timing of each Hallmark, serving as input for ASCETIC. The ASCETIC algorithm was then executed with a total of 100 resampling iterations to robustly estimate the timing of Hallmarks. The output of this analysis resulted in an evolutionary model representing the order of Hallmarks acquisition during cancer evolution.

## Acknowledgments

This research was funded by Cancer Research UK (C44767/A29458 and C23017/A27935) and EPSRC (EP/W007096/1).

