## Supplemental Figures for "The temporal evolution of cancer hallmarks"

**Supplementary figures**

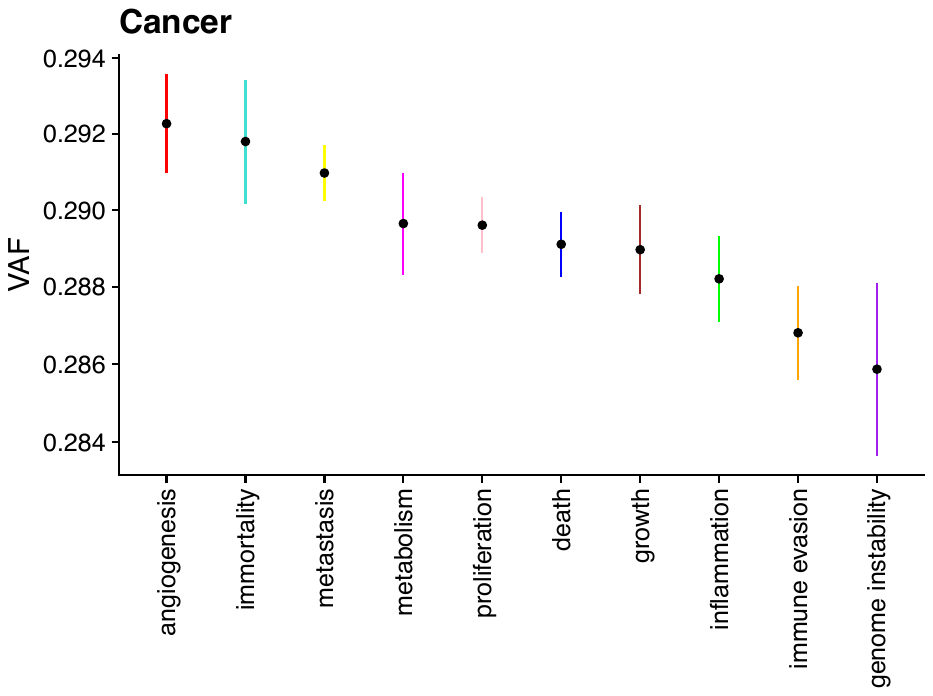

Supplementary Figure 1. Mean Variant Allele Frequency (VAF) for each cancer hallmark, derived from synonymous mutations only from all primary tumors in the TCGA database.

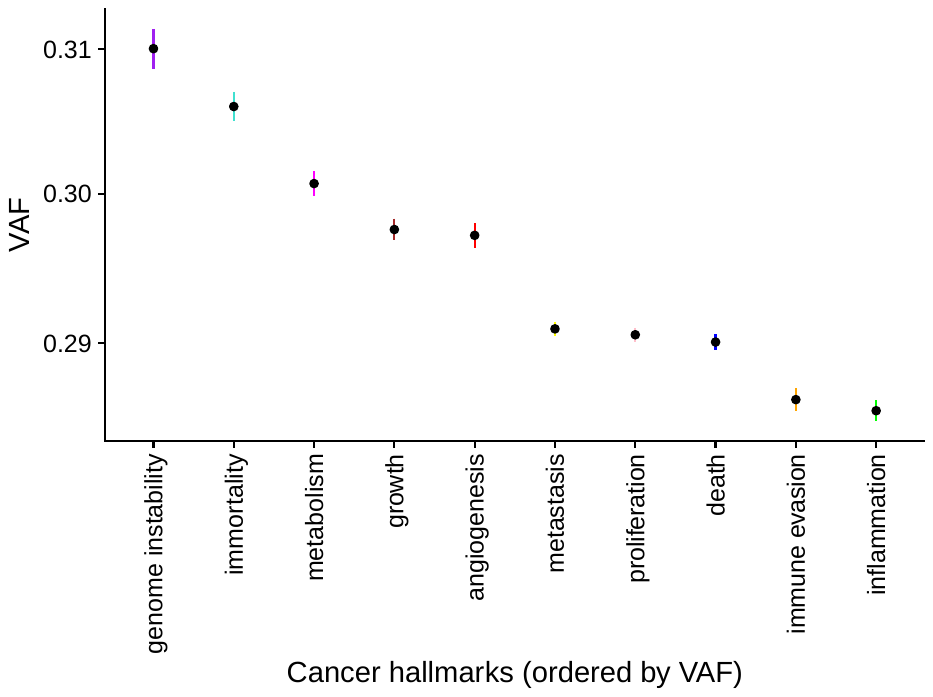

Supplementary Figure 2. Mean Variant Allele Frequency (VAF) for each cancer hallmark, derived from from all primary tumors in the TCGA database discarding tumors with evidence of aneuploidy (only ploidy 2 individuals).

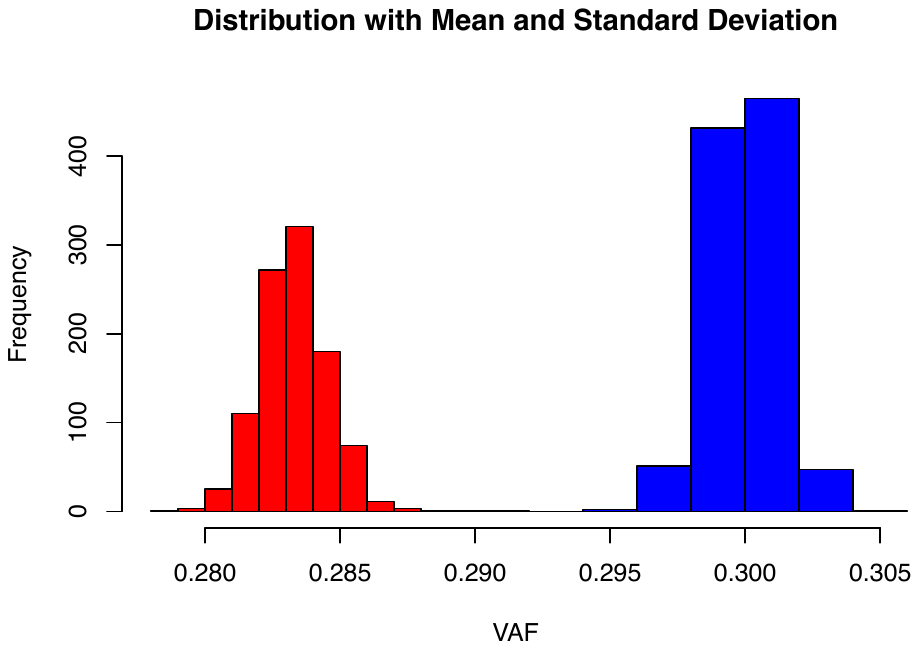

Supplementary Figure 3. Distribution of the mean VAF of random genes (neutral: red) and genes involved in one of the hallmarks (blue) across nonsynonymous mutations.

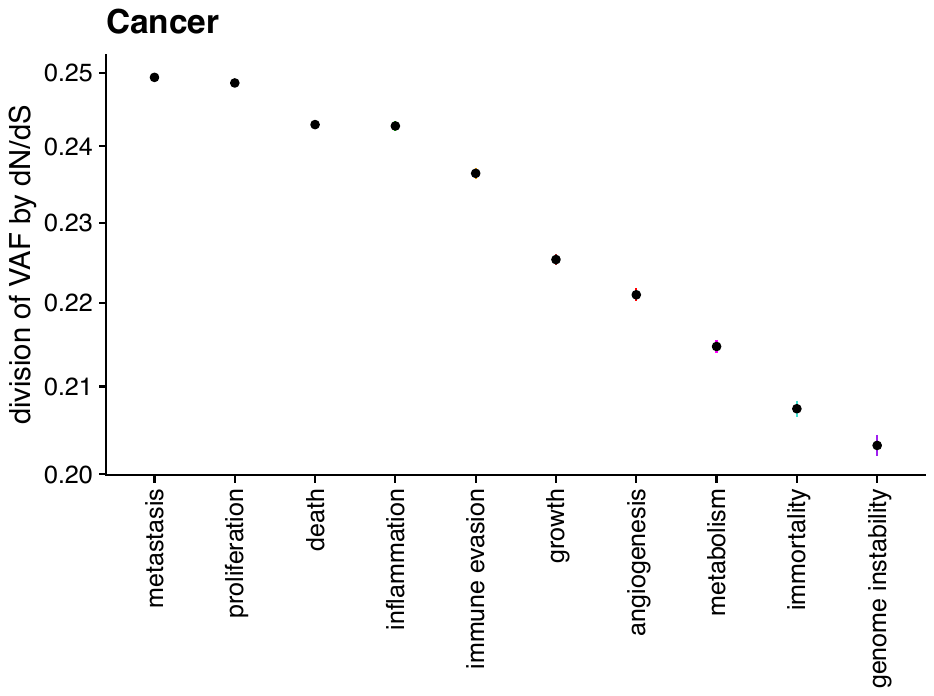

Supplementary Figure 4. Metric resulting from the division of VAF by dN/dS for each cancer hallmark from all primary tumors in the TCGA database.

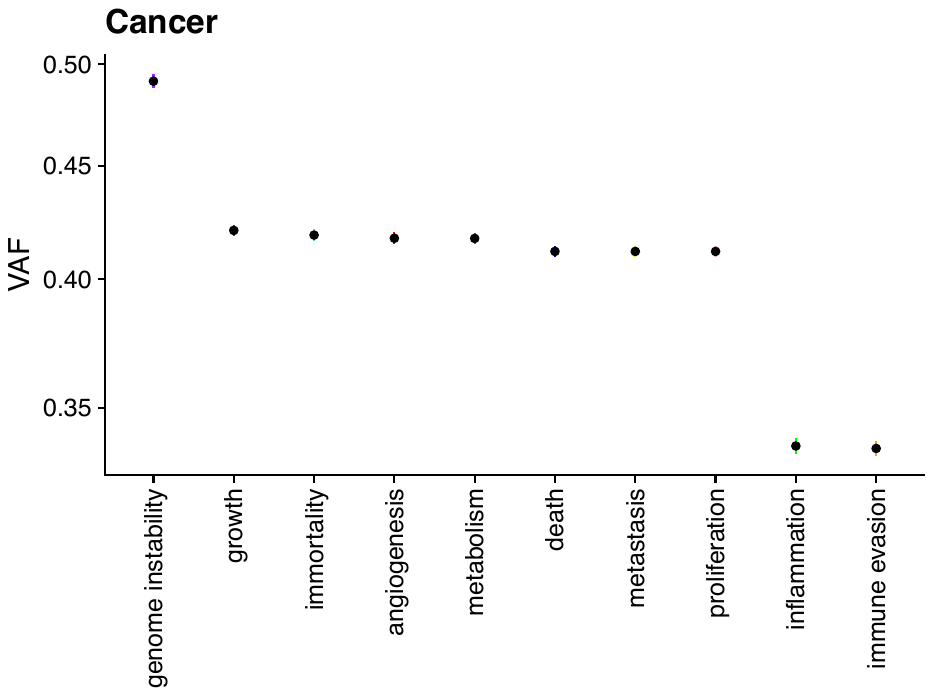

Supplementary figure 5. Mean Variant Allele Frequency (VAF) for each cancer hallmark, using only genes under positive selection from all primary tumors in the TCGA database.

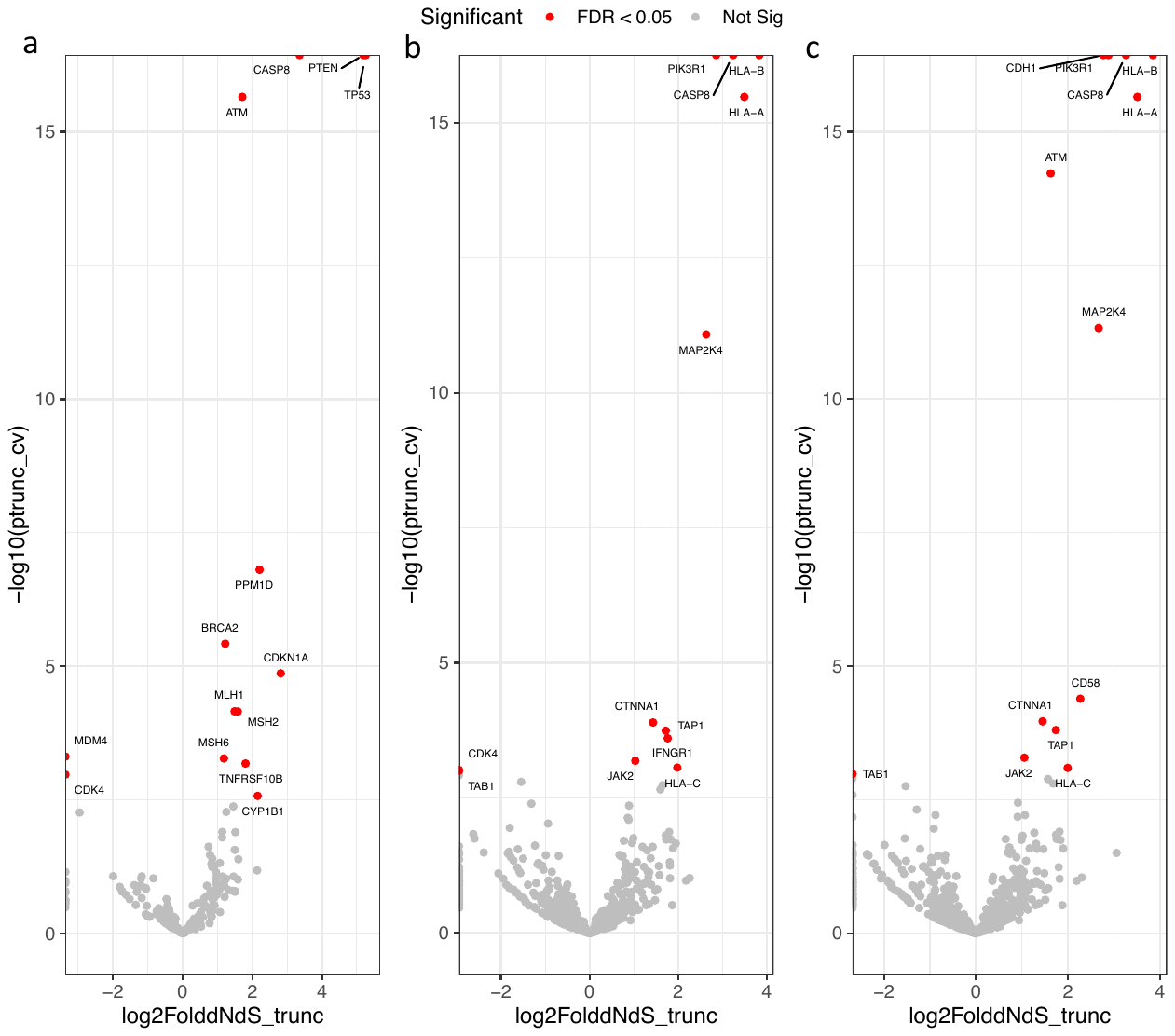

Supplementary figure 6. Volcano plot for dN/dS of nonsense mutations in a) genome instability genes, b) immune evasion genes, and c) inflammation genes.

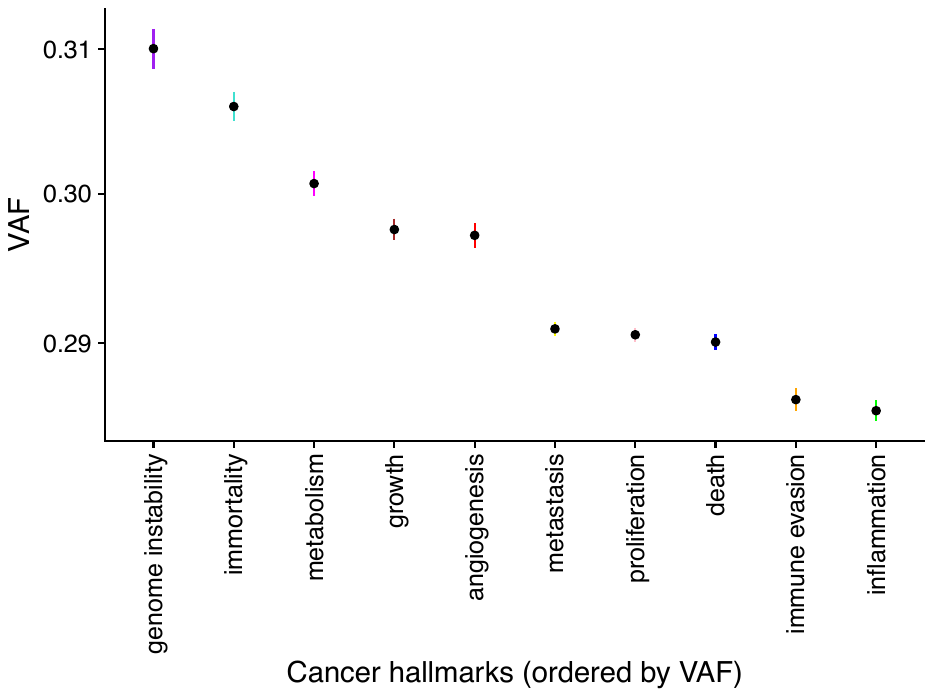

Supplementary Figure 7: Control for the VAF analysis by using copy number correction.

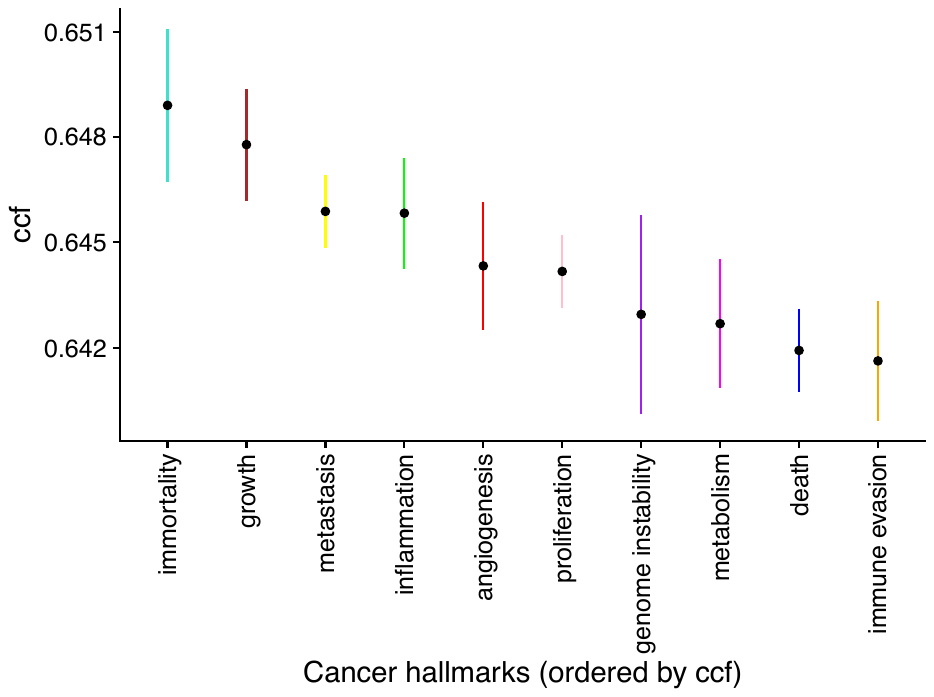

Supplementary Figure 8: Hallmark ordering analysis by using mean cancer cell fraction (CCF) instead of VAF.

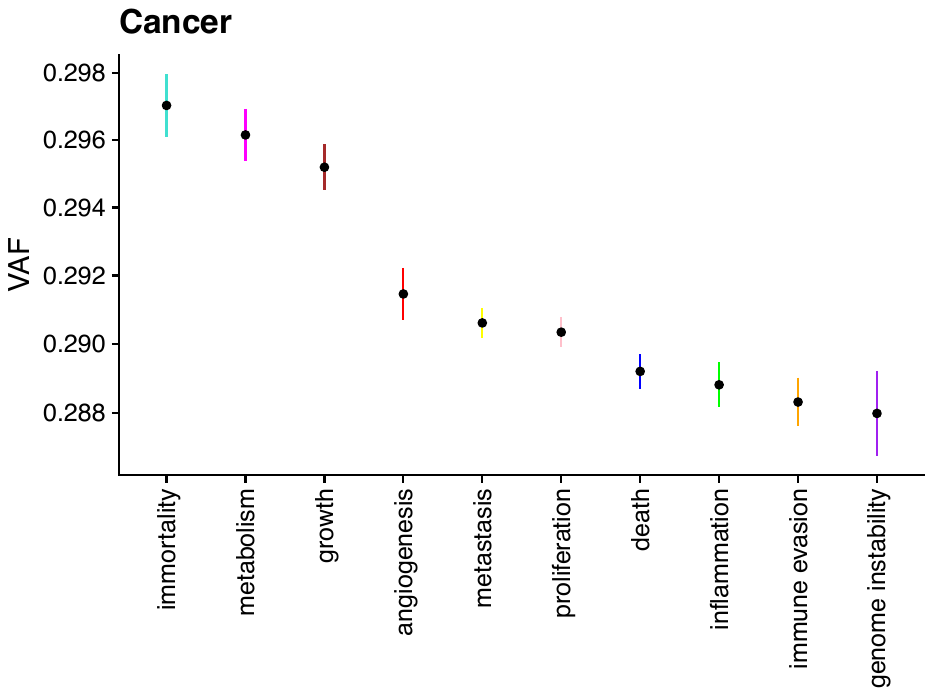

Supplementary Figure 9: Hallmark ordering of pancancer data without TP53 based on VAF.

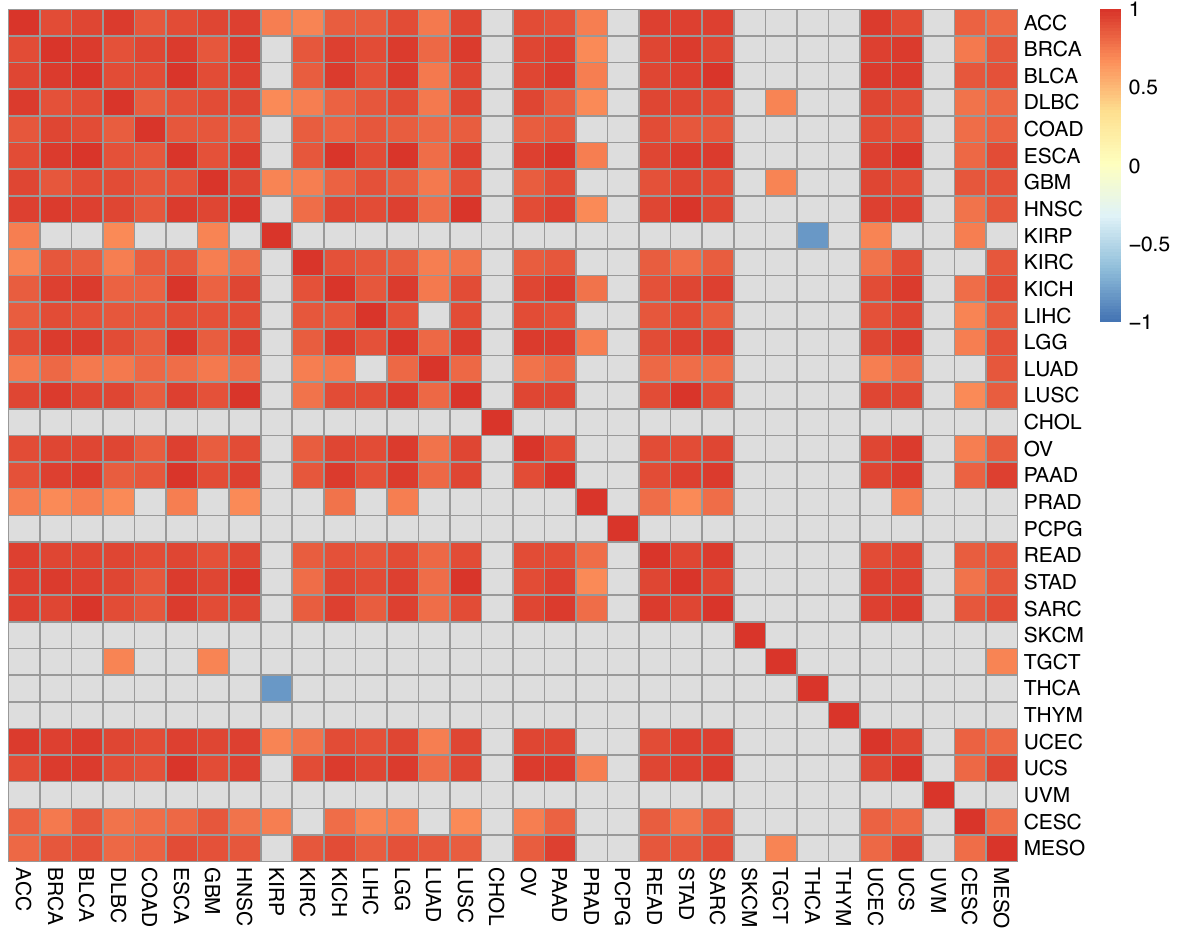

Supplementary Figure 10: Hierarchical clustering of spearman correlation values of the rank order according to VAF for 32 cancer subtypes. Colours are displayed only when the correlation is significant based on Benjamin Hochberg correction.

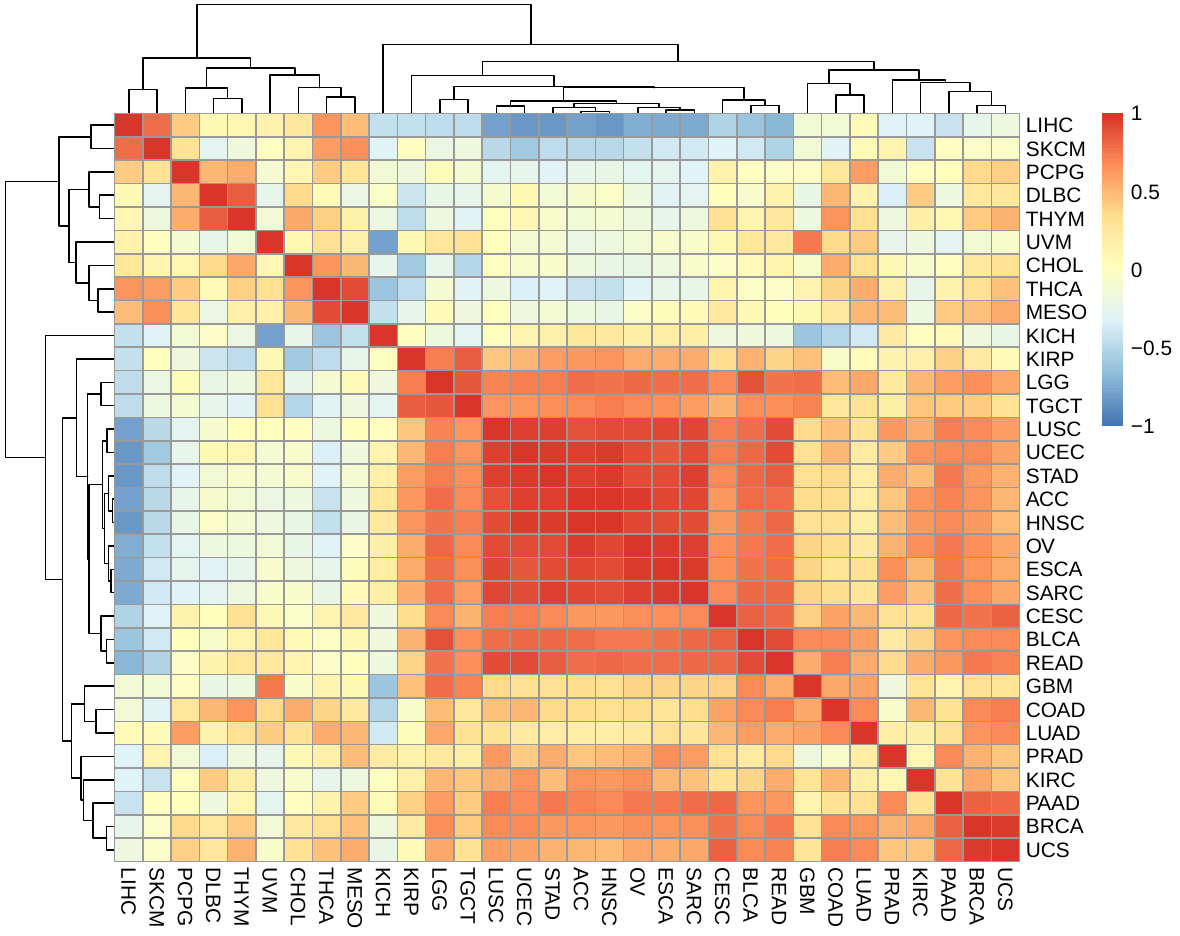

Supplementary Figure 11. Hierarchical clustering of spearman correlation values of the rank order according to CCF.

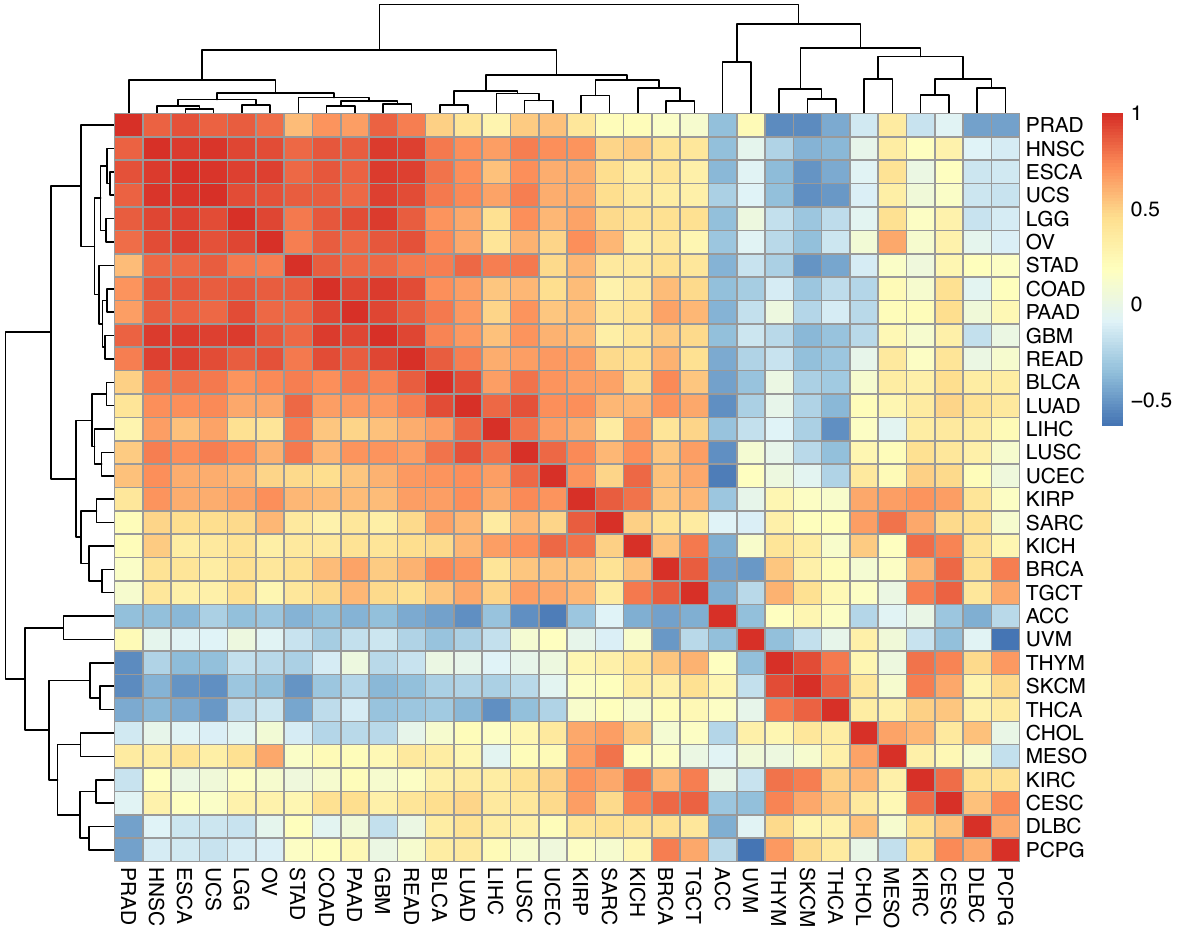

Supplementary Figure 12. Relative ordering of cancer hallmarks using Spearman correlation across 32 cancer types, according to dN/dS.

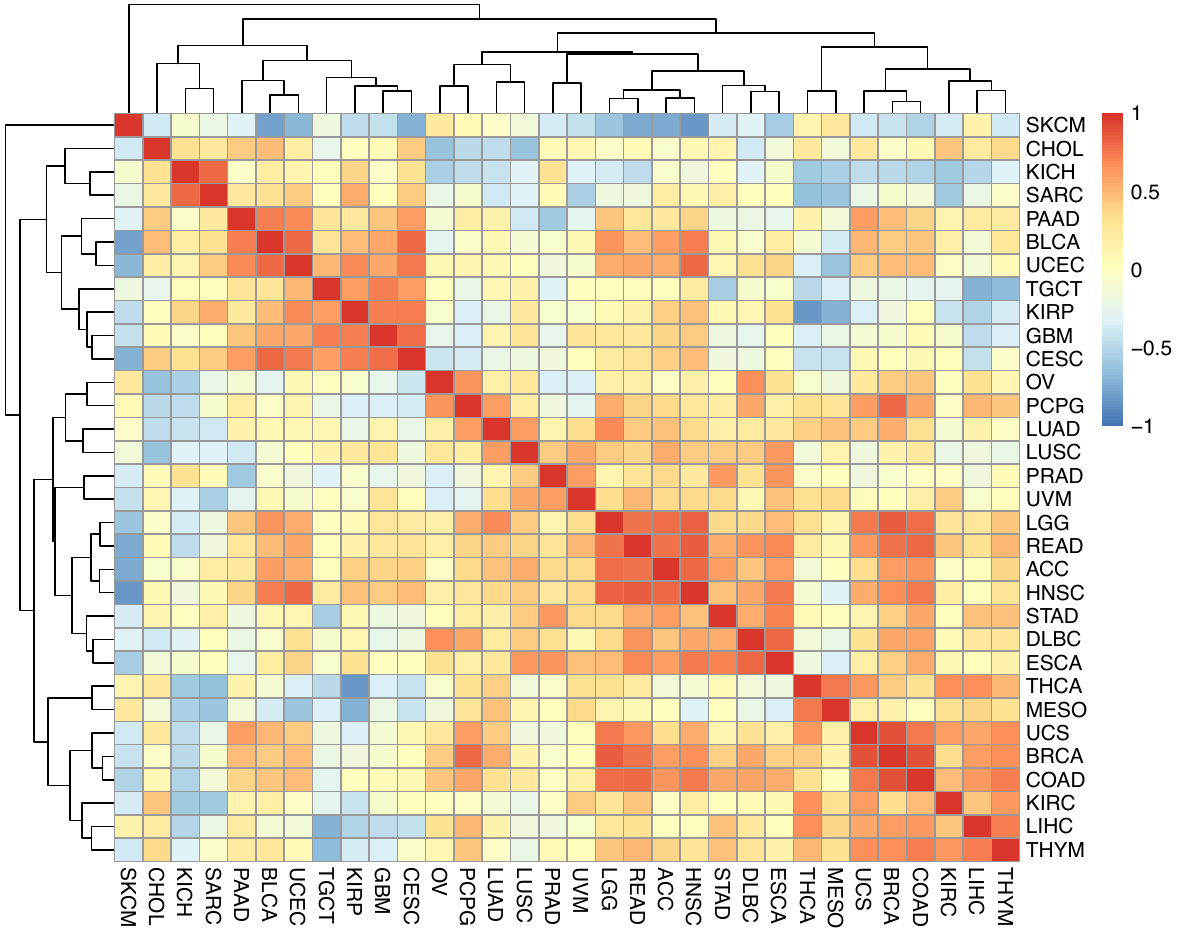

Supplementary Figure 13. Hierarchical clustering of spearman correlation values of the rank order according to VAF for 32 cancer subtypes without TP53. No correlation is significant based on Benjamin Hochberg correction.

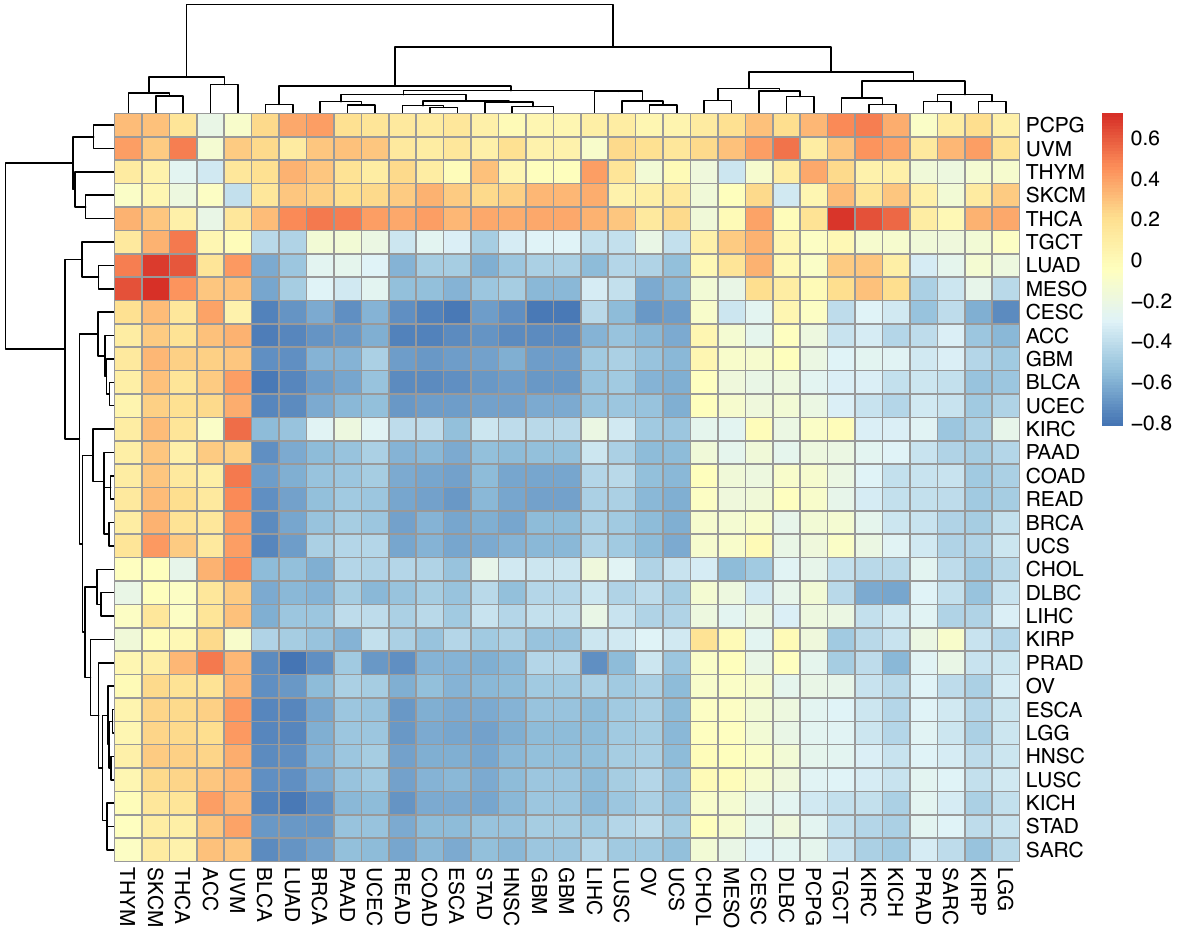

Supplementary Figure 14. Relative ordering of cancer hallmarks using Spearman correlation across 32 cancer types, comparing VAF and dN/dS ranking.

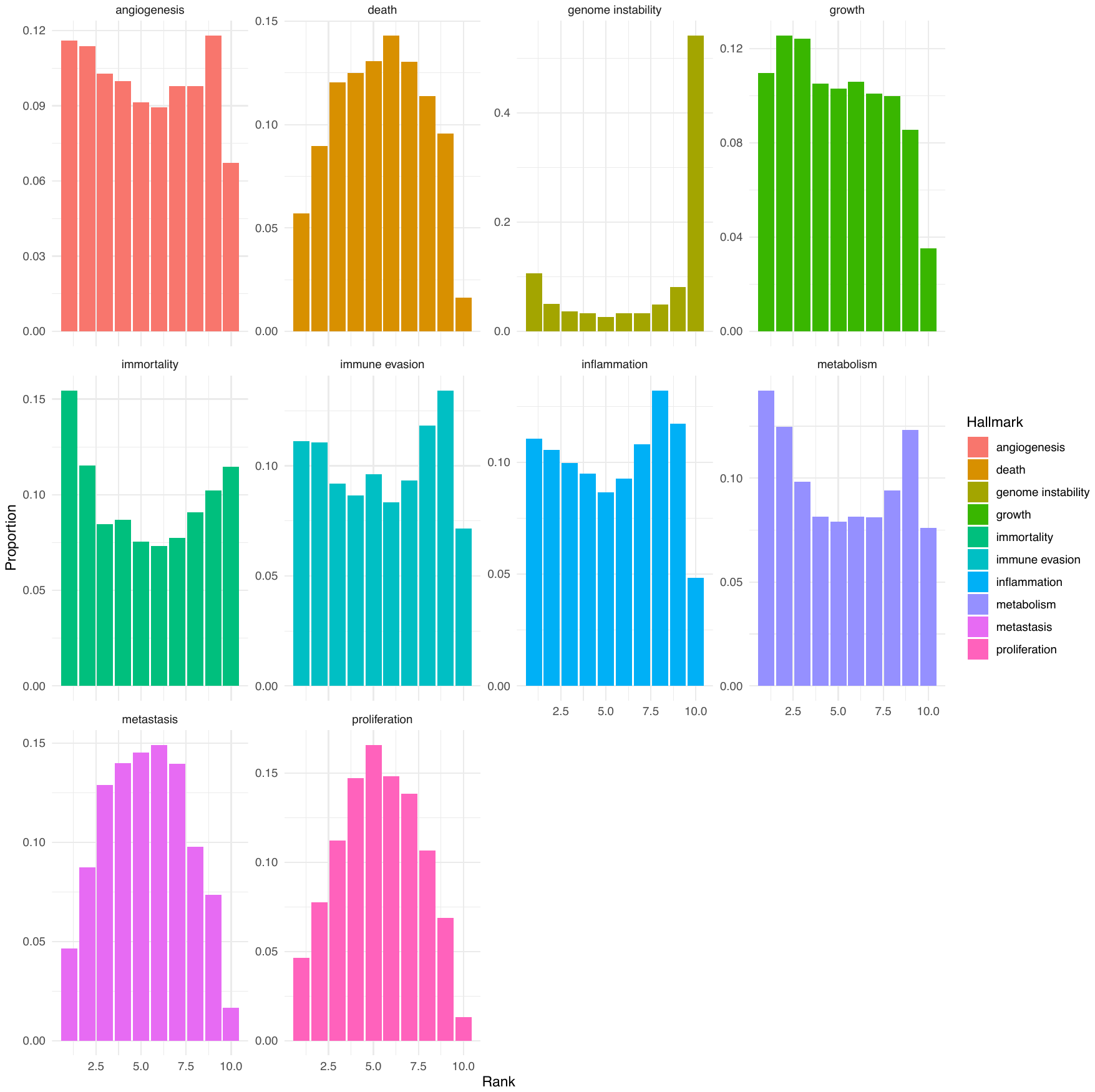

Supplementary Figure 15: Distribution of hallmark rank per patient from Pancancer data without TP53 based on VAF.

Supplementary Figure 16: Oncoprint plot showing the mutational landscape of genome instability genes.

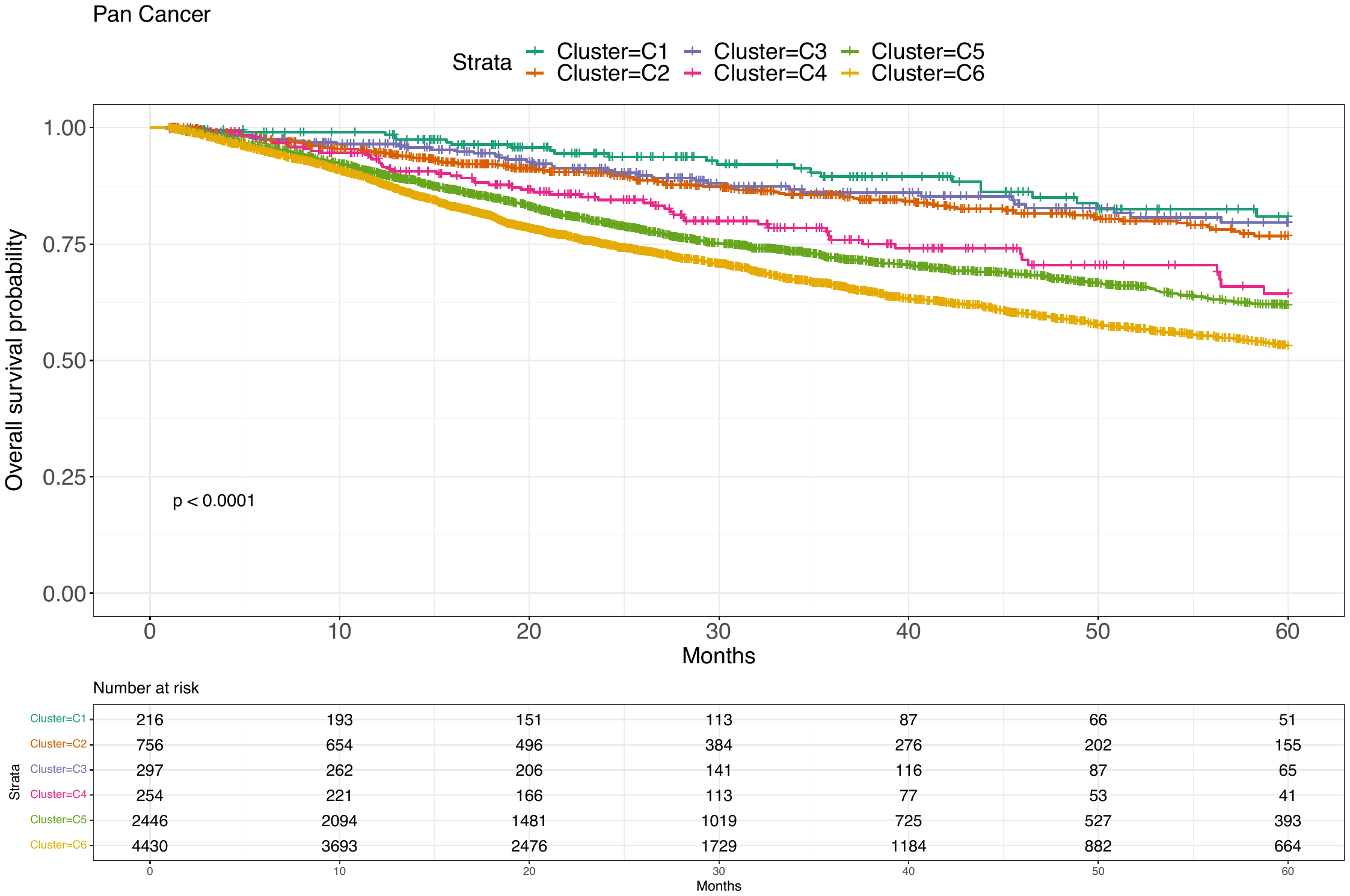

Supplementary Figure 17: Kaplan-Meier curves for 6 clusters predicted using ASCETIC algorithm in all patients from all tumor types.

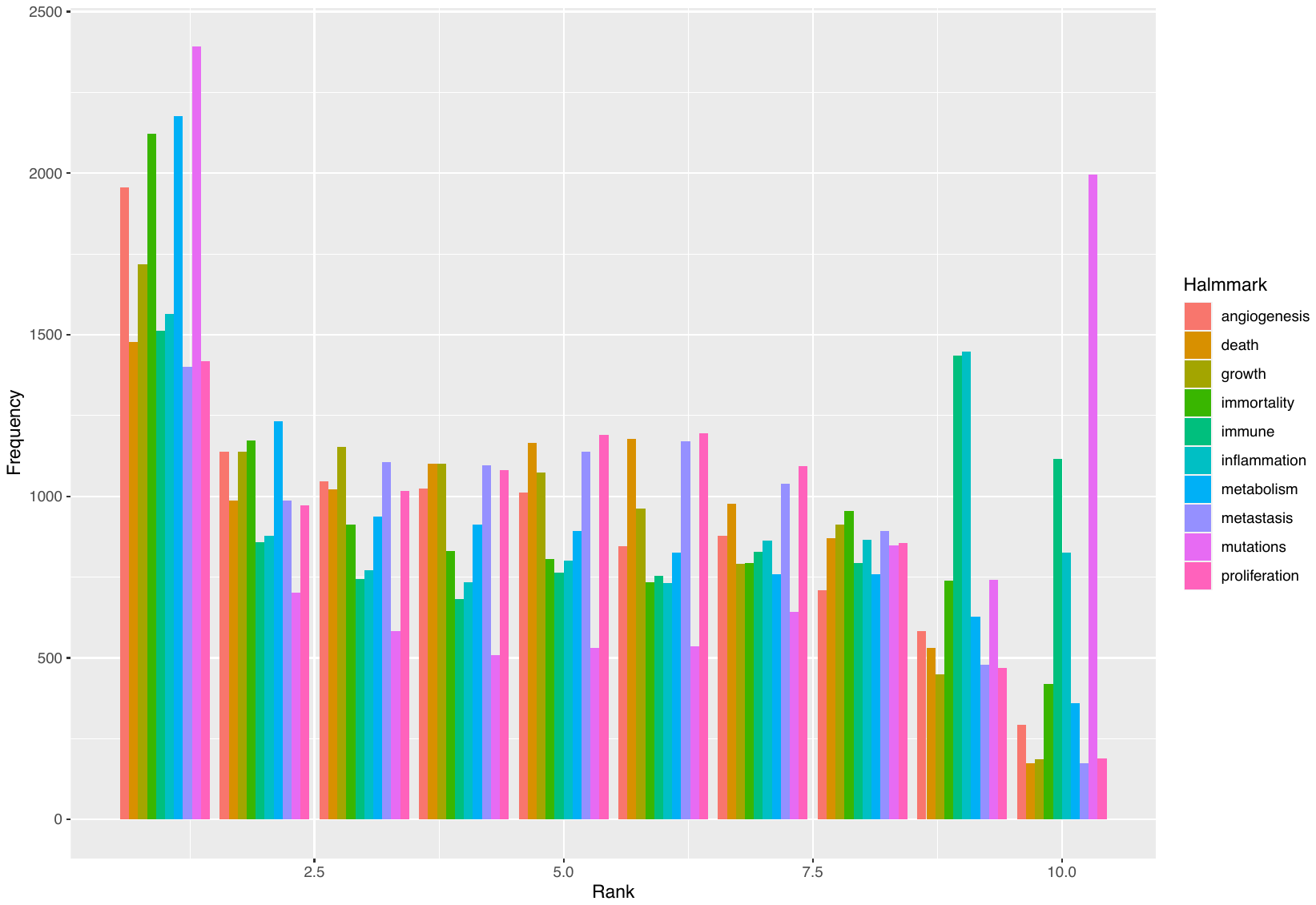

Supplementary Figure 18: Frequency of ranks for hallmarks.

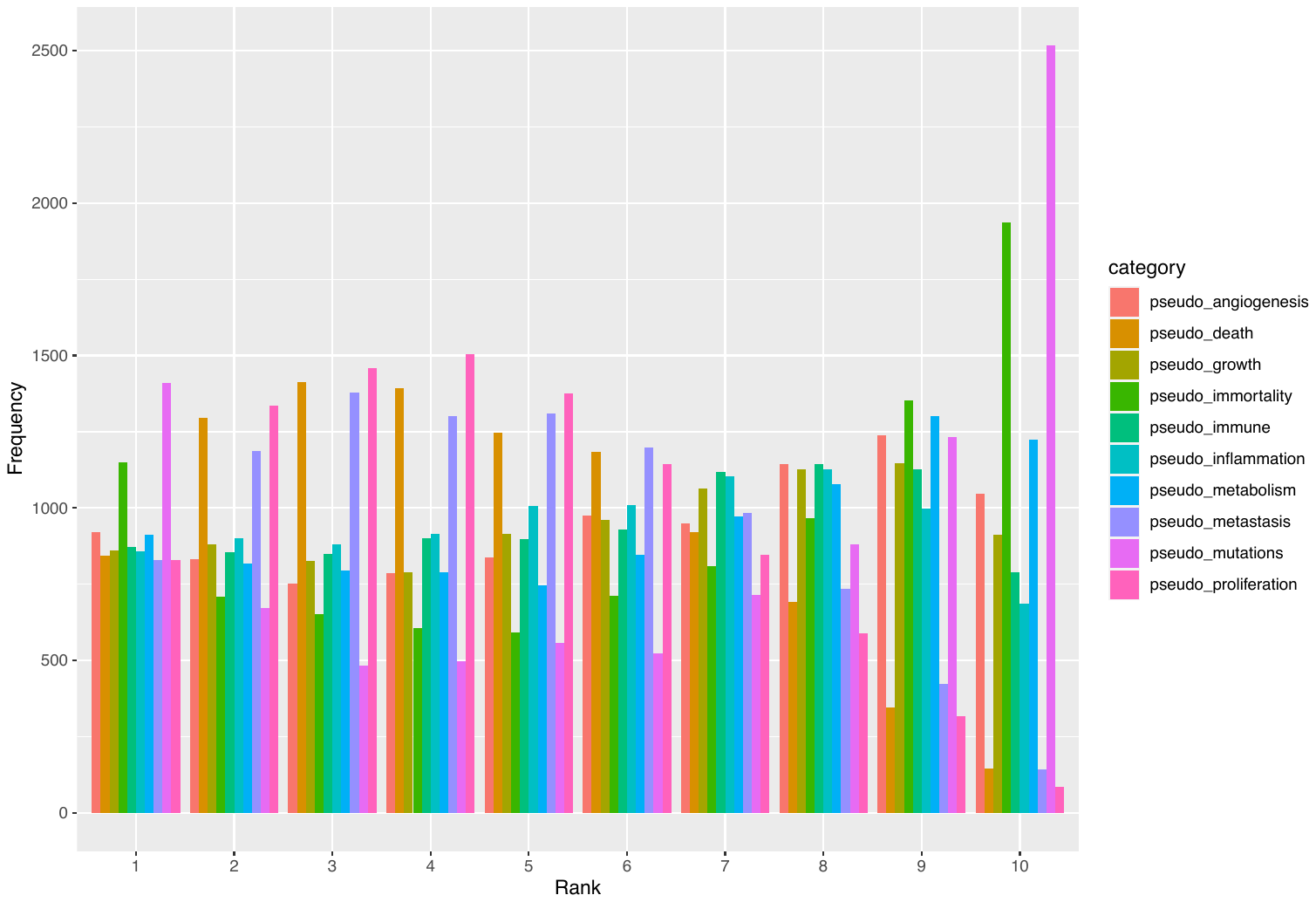

Supplementary Figure 19: Frequency of ranks for pseudohallmarks.
